# Bird assemblages in a peri-urban landscape in eastern India

**DOI:** 10.1101/2022.07.14.500013

**Authors:** Ratnesh Karjee, Himanshu Shekhar Palei, Abhijit Konwar, Anshuman Gogoi, Rabindra Kumar Mishra

**Affiliations:** PG Department of Wildlife and Biodiversity Conservation, North Orissa University, Takatpur, Baripada, Odisha, PIN - 757003, India; Aranya Foundation, Plot No-625/12, Mars Villa, Panchasakha Nagar, Dumduma, Bhubaneswar, Odisha, PIN - 751019, India

**Keywords:** Bird assemblage, diversity, composition, peri-urban landscape, feeding guild

## Abstract

Across the tropics, human settlements and agricultural lands have rapidly replaced the natural habitat; however, our understanding of the biodiversity value in such human-dominated landscapes is limited. To study the effects of heterogeneous habitats on bird diversity in the peri-urban landscape, we surveyed four different habitat types (residential areas, cropland and wasteland, water bodies, and sal forest) from February 2018 to January 2019. During the survey, 117 bird species were recorded belonging to 48 families and 98 genera in the study area. We observed that cropland and wasteland harboured rich avian diversity, among other sites. Both the Non-parametric Multidimensional Scale (NMDS) and one-way ANOVA Significant Test showed significant variations in bird species composition among habitats. Our study could be helpful in future conservation and management of the birds’ diversity to make peri-urban habitats more favorable.

## 1. Introduction

Birds are bipedal feathered organisms (Ali, 1990). They are ecosystem dwellers, in terrestrial and aquatic habitats with different food habits (Blair, 1999). Avifauna also acts as pollinators and seed dispersers and is closely associated with the food chain in the ecosystem (Nason, 1992; Niemi, 1985). Therefore, birds are important ecological indicators of any habitat (Blair, 1999; Niemi, 1985). Habitat destruction and human disturbances decreases avian species’ diversity and force them to inhibit in urban areas (Sarkar et al., 2009; Wang et al., 2022). In India, the wetlands are facing tremendous anthropogenic pressure (Prasad et al., 2002), which significantly impacts the structure of the avian community (Reginald et al., 2007; Verma et al., 2004), as they use seeds, plants, insects, other vertebrates or invertebrates as their diet (Hamel, 1982).

The overall community structure of birds in any landscape can be assessed by monitoring the species’ richness and abundance on a spatio-temporal scale (Sarkar et al., 2009). Such management practices may explain the role of environmental limiting parameters and anthropogenic factors’ interaction in determining the diversity and density of avifauna (Terborgh, 1985). Habitat loss and fragmentation due to anthropogenic pressures are primary drivers of global biodiversity decline (Garden et al., 2006; Sodhi et al., 2011). Forest fragmentation occurs when large, continuous forests are divided into smaller blocks by roads, clearing for agriculture, urbanization, and other human activities. Urban development for residential, commercial, and industrial properties’ was undeniably the most damaging, persistent, and rapidly expanding form of anthropogenic pressure (Miller and Hobbs, 2002; Vitousek et al., 1997).

There is a critical relationship between bird diversity and various environmental factors. Many studies discovered the mixture of bird diversity in various habitats like urban and rural habitats, farmland and forest habitats (Barth et al., 2015; Beukema et al., 2007; Fontana et al., 2011; Herzog et al., 2005; Shoffner et al., 2018; Tu et al., 2020). It is predicted that by the year 2050, most of the global population of birds will inhabit the urban landscape (Loss et al., 2009). Farmland, pastureland, and urban areas are important bird habitats as they hold much wildlife outside the protected areas (Beukema et al., 2007; Shoffner et al., 2018). Urban habitats encourage bird populations in cities and their surroundings; however, the urbanization process, like landscape conversion, is a great threat to the bird population (Wang et al., 2022). The growing human population and rapid landscape transformation for urban uses threaten biodiversity (Gatesire et al., 2014).

We studied bird assemblages to understand their abundance and distribution pattern in the heterogeneous landscape of Baripada, Odisha, India, a rapidly growing town blessed with agricultural land, forest areas, and several water bodies. We investigated four types of habitats: residential areas, cropland and wasteland, water bodies, and forest area to quantify the bird diversity and composition.

## 2. Materials and Methods

### 2.1. Study Area

Baripada was our study area which lay between 21.90°–21.96°N, 86.71°–86.78°E. It is located in the Chotanagpur Plateau Region of Odisha, Eastern India, with an altitude of 45m a.s.l (Fig. 1). It has a tropical climatic condition that experiences an extremely hot and dry summer followed by a humid monsoon and chilling winter with an annual temperature of 30° C. The winter season is observed between December to February. After that, summer continues from March to May, followed by the monsoon season from June to October, with an annual mean rainfall of 1,800 mm. Our study focused on Baripada and its outskirts surrounded by scrublands and agricultural lands with human interfered (Das et al., 2021).

**Fig. 1.**
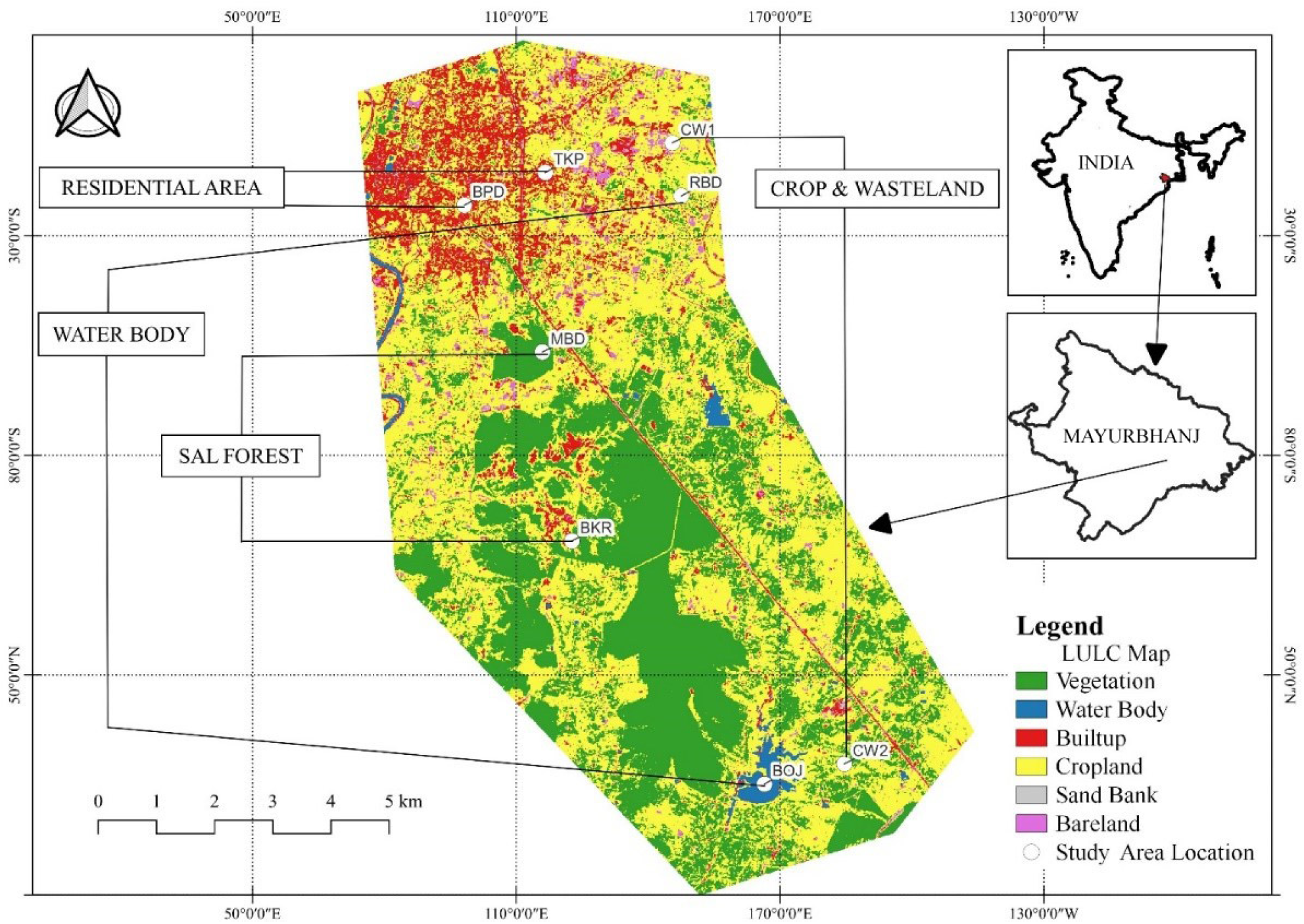
Map showing the different habitats and transect location in the city of Baripada, Odisha, India.

We selected four habitat types for bird sampling in a peri-urban landscape of Baripada, two transects in residential areas (RA), two in water bodies (WB), two in Sal (*Shorea robusta*) forest habitats (SF), and two in mixed habitats of cropland and wasteland (CW) (Fig. 1, Table 1).

**Table 1.** Description of different habitat types in the study area.

| Habitat | Transect* | GPS Coordinate |  | Characteristics of the study area |
| --- | --- | --- | --- | --- |
|  |  | Lat (N) | Lon (E) |  |
| RA | BPD | 21.928153 | 86.737023 | Human dominated landscape with less vegetation covers |
|  | TKP | 21.933584 | 86.750474 | Human dominated landscape with vegetation covers |
| WB | RBD | 21.929695 | 86.773065 | Small pond with aquatic flora and medium patches of vegetation covers |
|  | BOJ | 21.831909 | 86.786870 | Large pond with aquatic flora and small vegetation covers |
| CW | CW1 | 21.938516 | 86.771574 | Mosaic landscape of annual crops and wasteland or fellow land with small patches of vegetation, grass, seasonal canals and ditches. |
|  | CW2 | 21.835280 | 86.800143 | Mosaic landscape of annual crops and wasteland or fellow land with small patches of vegetation, grass, seasonal ditches |
| SF | MBD | 21.903739 | 86.749982 | Sal dominated forest covers with small canal and open area |
|  | BKR | 21.872335 | 86.754893 | Sal dominated forest covers with small canal and open area |

\*BPD = Baripada, TKP = Takatpur, RBD = Rani Bandh, BOJ = Borjhor Dam, CW1 = Crop & Waste land near University, CW2 = Crop & Wasteland near Borjhor, MBD = Monchabandha Forest, BKR = Bhudikhamari Forest

### 2.2. Field survey and avifaunal sampling

We studied bird diversity and abundance from February 2018 to January 2019 in the study area using point counts set along line transects (Bibby et al., 2000), following an approach comparable to that used in other large-scale bird surveys (Sauer et al., 2013). A total of 8 transects were established, with 18 replications in each transect to obtain a spatially homogenous distribution. Transects were about 5 km (0.62 mean; SE ± 0.9 km, 0.66 range) long. They were demarcated before surveys using 1:25,000 topo maps, aiming to obtain a sufficient length route linear and continuous as possible. In each transect, the first sampling point coincided with the beginning of the transect; all the other points were set 200 m apart. This spacing was considered sufficient to avoid double counts (Dias et al., 2013). At each sampling point birds were recorded from a fixed raising position within a circle of 50 m radius for a specific period (10 min) at every point (Camacho-Cervantes et al., 2018; Issa, 2019; Ralph et al., 1996). The counting of avian species was conducted during bird’s peak activity (up to 3 hours after sunrise), in the early morning after the sun rises (Camacho-Cervantes et al., 2018; Ralph et al., 1996). A minimum of one or two counts was established in each transect every month in each site.

### 2.3. Bird identification, taxonomy and nomenclature

A binocular (Nikon 10 × 40) and a camera (Canon S 325) were used to identify and document encountered bird species. The photographic evidence was again verified using a reference field guidebook (Grimmett et al., 2016) to avoid biases and identify taxonomic issues of avian species. Overflying birds were not included, as they could be only moving through or above the surveyed habitat. Bird sampling limited to those individuals who had evidence to use the habitat was included in the study.

### 2.4. Nonparametric richness estimation

We plotted a species accumulation curve to evaluate whether the number of bird species sampled was representative of the bird community. IndividualLbased rarefaction curves were used to compare species richness at the habitat level. Nonparametric species richness estimators such as Chao 1, Jackknife 1, and Bootstrap were calculated for different habitats. Nonparametric species richness estimators like Chao 1 index were selected because of their ability to deal with uneven and small sampling sizes (Chao, 1984). Jackknife 1 allows the estimated richness for bias reduction (Walther and Moore, 2005).

### 2.5. Species richness and diversity

Species richness is estimated as the number of bird species present in a particular habitat. Shannon’s diversity index (*H*□) was calculated by multiplying the proportion of each species by their natural log. Shannon Index (*H*□) = □*pi*log(*ln*)*pi*, where *pi* is the proportion (*n/N*) of individuals of a particular species found (*n*) divided by the total number of individuals recorded (*N*), *ln* is the natural log, and □ is the sum of the calculations. Similarly, to understand the dominated species within the community, Simpson diversity index (*D*) was calculated by using the formula as Simpson Index (*D*) = 1-□n (n-1)/*N* (*N*-1), where n is the total number of birds of a particular species and N is the total number of birds of all species. The evenness of bird species compares the similarity of the population size of each species. Evenness Index (*J*′) was calculated using the ratio of observed diversity to maximum diversity using the equation (Kiros et al., 2018). Evenness Index (*J*′) = *H*□/*H*_max_, where *H*′ is the Shannon Wiener Diversity index and *H*_max_ is the natural log of the total number of species. Rank abundance plots were constructed to investigate species abundance distributions between habitats.

### 2.6. Bird assemblage and similarity

Two similarity indices were calculated to estimate shared species richness between habitats and different seasons. These included qualitative similarity estimates using the Jaccard index and Morisita-Horn index (Chao et al., 2005; Magurran, 2004). Besides, bird richness and diversity similarity were determined using Bray-Curtis similarity or distance index, which is formulated as BC_*ij*_*=* 1-(2C_*ij*_ / S_*i*_ + S_*j*_) where *i* and *j* are the two habitats, S_*i*_ and S_*j*_ are the total numbers of birds counted on *i*th and *j*th and C_*ij*_ is the only lesser count for each bird species counted in both habitats. In the Bray-Curtis similarity index, a value nearer to 0 means the communities have the same species composition, and a value closer to 1 means no share of any species.

To examine changes in bird functional diversity among different habitats, we classified birds into various guilds based on their diets: carnivorous (C), frugivorous (F), granivorous (G), omnivorous (O), insectivorous (I), and nectarivorous (N). A heat map has been produced to understand the spatio-temporal assemblage of birds’ feeding guild. Although Indian birds have mixed food habits, a simplified food guild based on the predominant food habits of each bird species was followed in this study. The classification of feeding guilds is followed by Grimmett et al., (2016) and Sengupta et al., (2014).

### 2.7. Statistical analysis

The non-parametric multidimensional scale (NMDS) test was performed to check the significant variation of bird community amongst the habitats using the permutation test (999 permutations). After that, a one-way ANOVA test was run to check the variation of birds’ richness, diversity, and abundance in the study area. Again, a multiple comparison Tukey’s test was performed to quantify variation among the habitats. The statistical analyses were done in R 4.0.2 statistical data processing packages. Species accumulation curves, diversity indices, rank abundance plots, habitat share Venn diagram, and heatmap were calculated using “BiodiversityR”, “VennDiagram”, and “superheat” packages.

## 3. Results

### 3.1. Species richness and diversity

During the survey, 117 bird species were recorded belonging to 48 families and 98 genera within the four different habitats (Appendix A). Of these, 85.83% are the resident species (103 bird species), while 9.17% (11 bird species) are winter migrants. The highest bird richness was observed in CW (64 species; evenness J=0.91), followed by RA (56 species; evenness J=0.90) and SF (54 species; evenness J=0.88). The WB represents the lowest species richness (37 species; evenness J=0.78). The individual-based rarefied richness curve of bird species reached an asymptote for all habitats, indicating that sampling effort was sufficient (Fig. 2). The maximum value of Shannon-Wiener Index was recorded in CW (H□=3.79), followed by RA (H□=3.63), SF (H□=3.51), and lowest in WB (H□=2.84). The value of Simpson’s index of CW scored highest (D=0.97), followed by RA (D=0.96), SF (D=0.95), and WB (D=0.90).

**Fig. 2.**
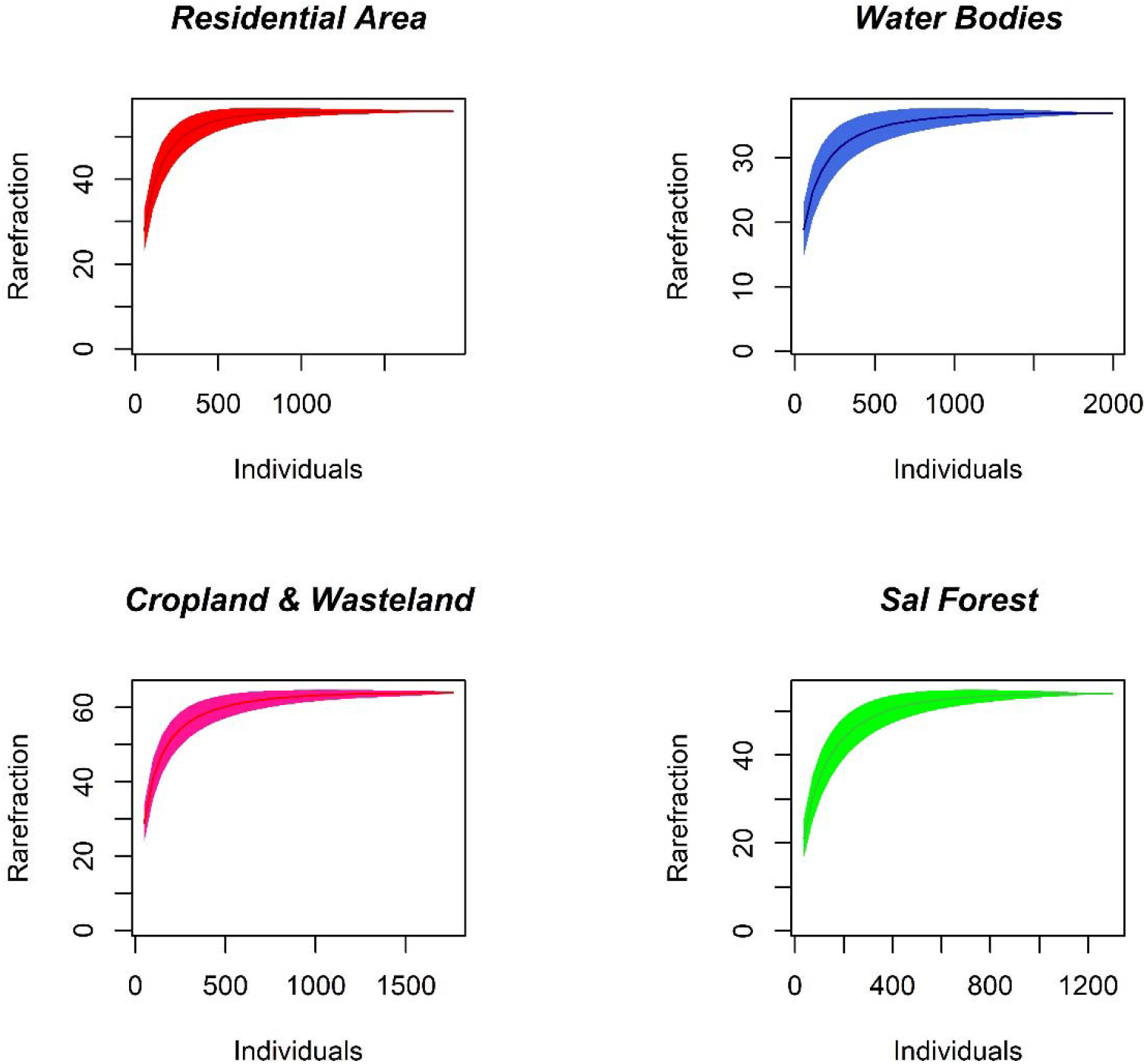
Individual based rarefaction curve for bird species richness found in four different habitats in the study area. Shaded area around the curve indicates 95% confidence intervals (*CI*).

The NMDS results (non-metric fit, R^2^ = .0.936, linear fit, R^2^ = 0.695, stress = 0.252) showed that bird community were significantly varied (MAST, F=17.50, DF=3, R^2^ =0.272, p =0.001) (Fig. 3). In addition, one-way ANOVA suggested that there is a significant variation in bird richness (F=13.70, DF=3, P<0.001) and abundance (F=6.61, DF=3, P<0.001) among habitats (Fig. 4). Similarly, Tukey’s HSD test for multiple comparisons also showed significant variation in bird richness only between SF and CW (P<0.05, 95% C.I. =[−5.22, −0.003]);WB and CW (P <0.001, 95% C.I. = [−8.27, −3.05]); WB and RA (P<0.001, 95%, C.I. = [−7.97, −2.75]); WB and SF (P<0.01, 95%, C.I. = [−5.66, −0.44]). On the other hand the mean value showed significant difference in abundance only between SF and CW (P<0.05, 95%, C.I. = [−25.59, −0.85]); SF and RA (P<0.05, 95%, C.I. = [−28.93, −14.18]); WB and SF (P<0.001, 95%, C.I. = [7.30, 32.03]).

**Fig. 3.**
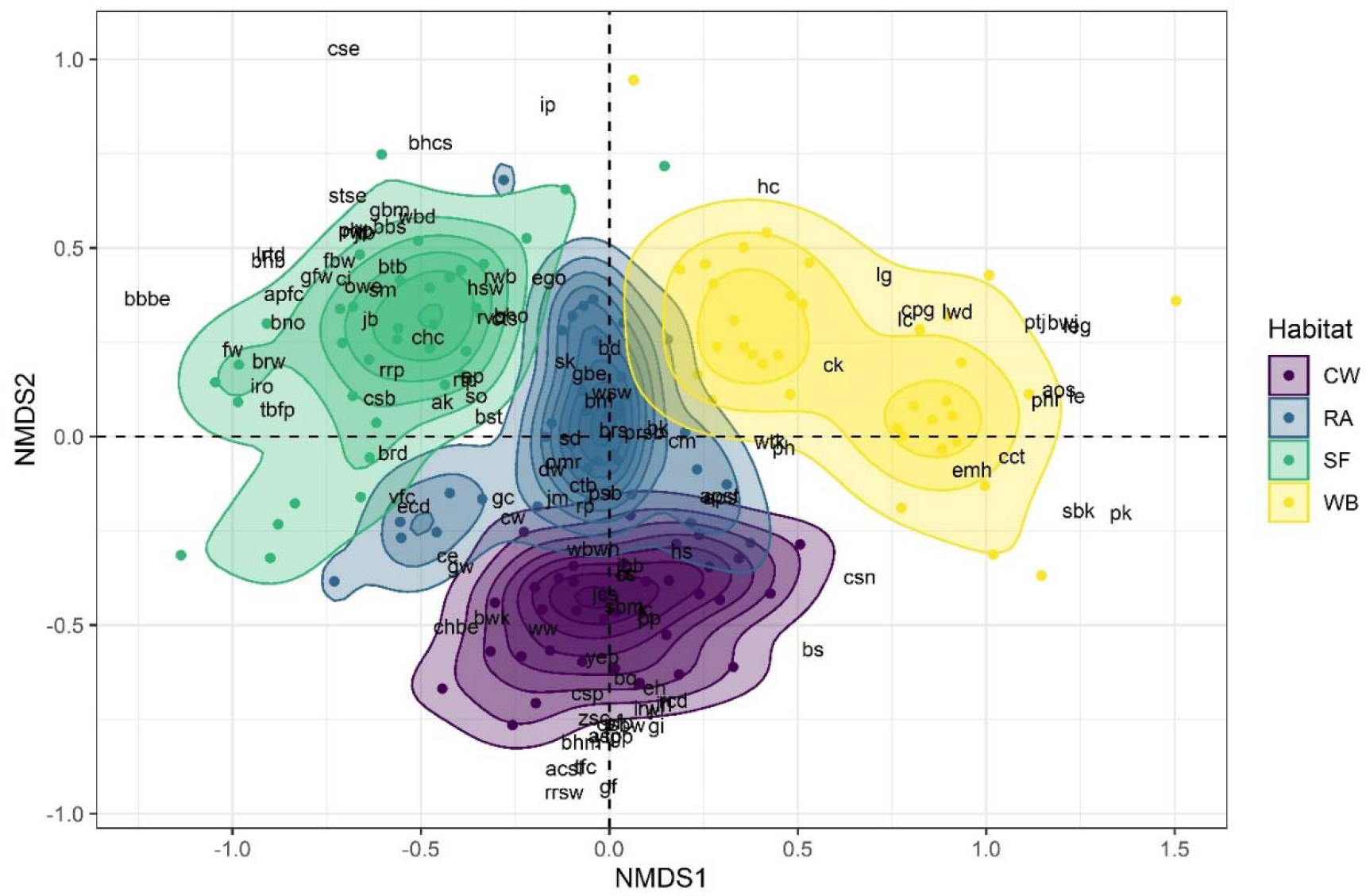
Non-metric Multidimensional Scaling showing the different bird species composition at each habitat (stress = 0.25; Non-metric fit R^2^ = 0.936; linear fit R^2^ = 0.695); CW= Crop and Wasteland, RA= Residential habitat, SF= Sal Forest and WB= Water Body.

**Fig. 4.**
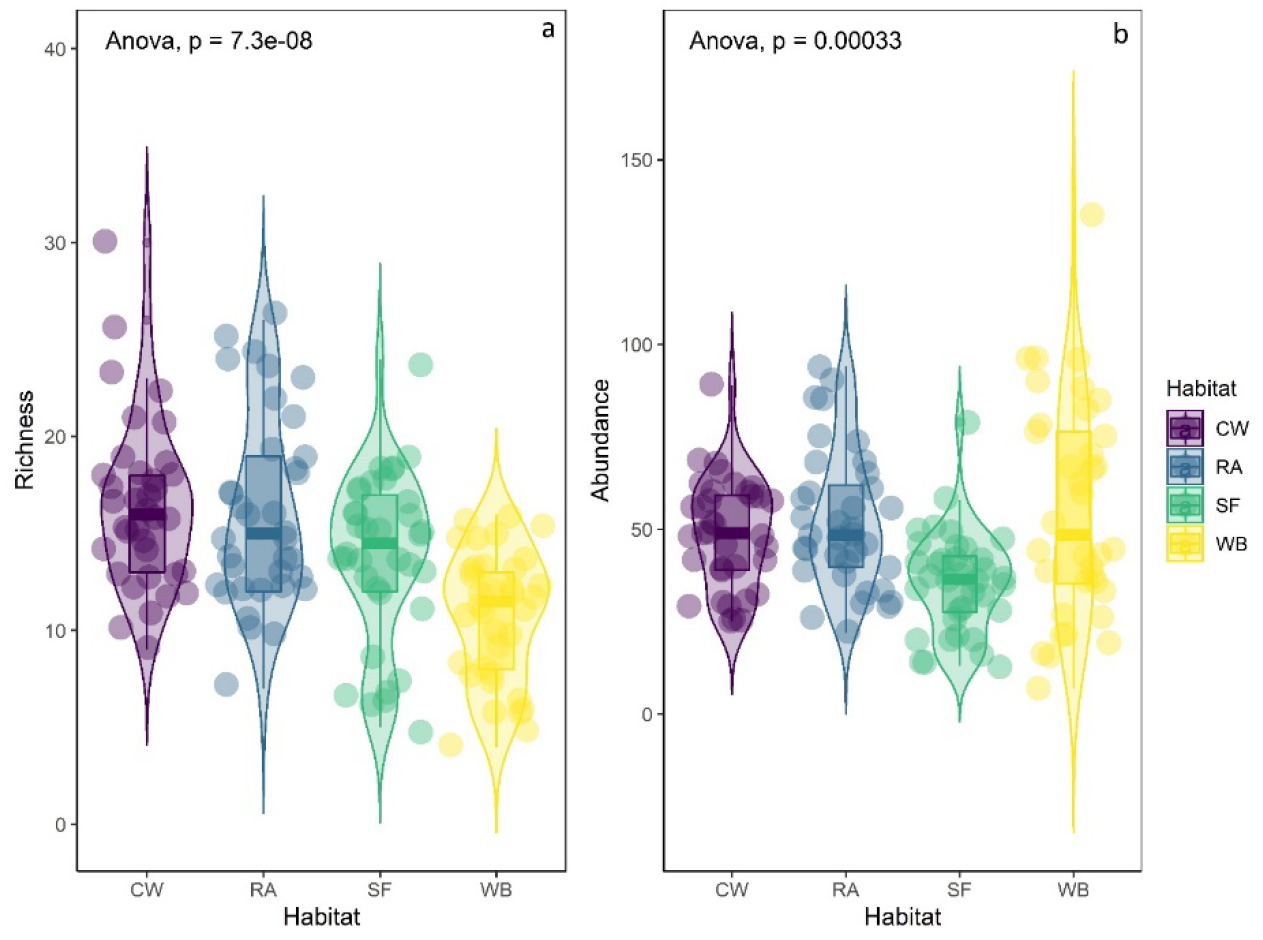
ANOVA showing that there was a significant differences between habitat in terms of bird richness and abundance; CW= Crop and Wasteland, RA= Residential habitat, SF= Sal Forest and WB= Water Body.

### 3.2. Bird rank-abundance

A total of 6963 individual birds were counted, including 1992 in WB (28.61%), 1912 in RA (27.46%), 1760 in CW (25.28%), and 1299 in SF (18.66%).. Overall taxa level analysis showed that *Dendrocygna javanica* represented the highest individual (n=461), followed by *Columba livia* (n=191), *Bubulus ibis* (n=147), and *Sturnia malabarica* (n=121). In WB, *Dendrocygna javanica* (n=461) is the dominating rank over *Tachybaptus ruficollis* (n= 282), *Nettapus coromandelianus* (n=166), *Microcarbo niger* (n=136) and *Anastomus oscitans* (n=101). *Columba livia* is dominating rank-abundance bird in RA, including 191 individuals, followed by *Bubulcus ibis* (118 individuals), *Sturnia malabarica* (117 individuals), *Gracupica contra* (82 individuals), and *Merops orientalis* (81 individuals). In CW, the rank-abundance plot showed that *Bubulcus ibis* (147 individuals) is placed at the top, followed by *Gracupica contra* (103 individuals), *Columba livia* (90 individuals), *Acridotheres tristis* (76 individuals), and *Cypsiurus balasiensis* (74 individuals). In SF rank-abundance, *Sturnia malabarica* was dominating with 121 individuals over the rest of the species. In addition, *Pycnonotus cafer* and *Bubulcus ibis* placed in the second and third rank with 98 and 96 individuals, followed by *Psittacula krameri* and *Merops orentalis* with 75 and 66 individuals, respectively (Fig. 5).

**Fig. 5.**
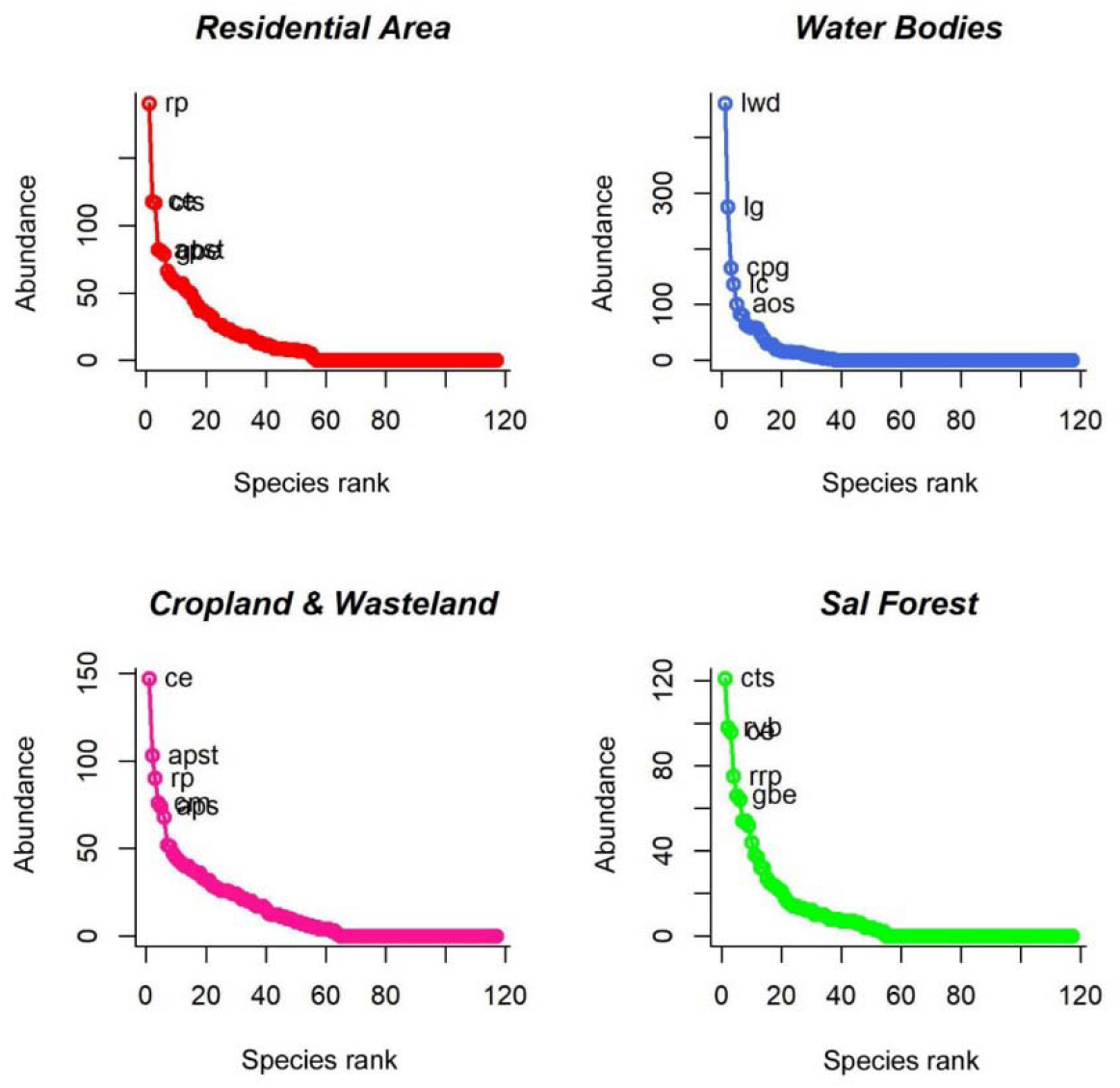
The rank-abundance representing the position of the bird in various habitats; the highest abundance of each birds in each habitat held the top position. Please see Appendix A for abbebaration of each birds

### 3.3. Similarity and shared species richness

In our study, we recorded nine bird species *Gracupica contra, Dicrurus macrocercus, Acridotheres tristis, Motacilla flava, Merops orientalis, Pycnonotus cafer, Pycnonotus jocosus, Accipiter badius*, and *Spilopelia chinensis* which are found in all habitats. The highest bird richness was shared between RA and CW with a total of 39 species, followed by CW and SF with 25 species. SF and WB shared very low bird species (S=13) (Fig. 6). Bray-Curtis and Jaccard Indices suggested that there were higher dissimilarities between WB and SF (BC=0.87) and higher similarities between RA and CW (JI=0.52). Morisita-Horn Index also revealed that the compositional similarity of birds is higher between RA and CW (MH=0.27), followed by RA and WB (MH=0.91), and SF and WB (MH=0.93). The Euclidian distance index also suggested a higher similarity between RA and CW (EU=256), followed by CW and SF (EU=258). In contrast, RA and WB exhibited higher dissimilarity (EU=687), followed by WB and SF (Table 2).

**Table 2.** Pairwise comparisons of the bird communities among four different habitat types in the study area.

| Simillarity Index | Euclidian Distance |  |  | Bray-Curtis |  |  | Morista-Horn |  |  | Jaccard |  |  |
| --- | --- | --- | --- | --- | --- | --- | --- | --- | --- | --- | --- | --- |
|  | RA | WB | CW | RA | WB | CW | RA | WB | CW | RA | WB | CW |
| WB | 687.18 |  |  | 0.81 |  |  | 0.91 |  |  | 0.67 |  |  |
| CW | 255.83 | 653.68 |  | 0.47 | 0.78 |  | 0.27 | 0.89 |  | 0.52 | 0.77 |  |
| SF | 305.01 | 656.31 | 281.30 | 0.54 | 0.87 | 0.66 | 0.43 | 0.93 | 0.48 | 0.69 | 0.83 | 0.81 |

\*CW: Cropland and Wasteland, RA: Residential Areas, SF: Sal Forest, WB: Waterbodies

**Fig. 6.**
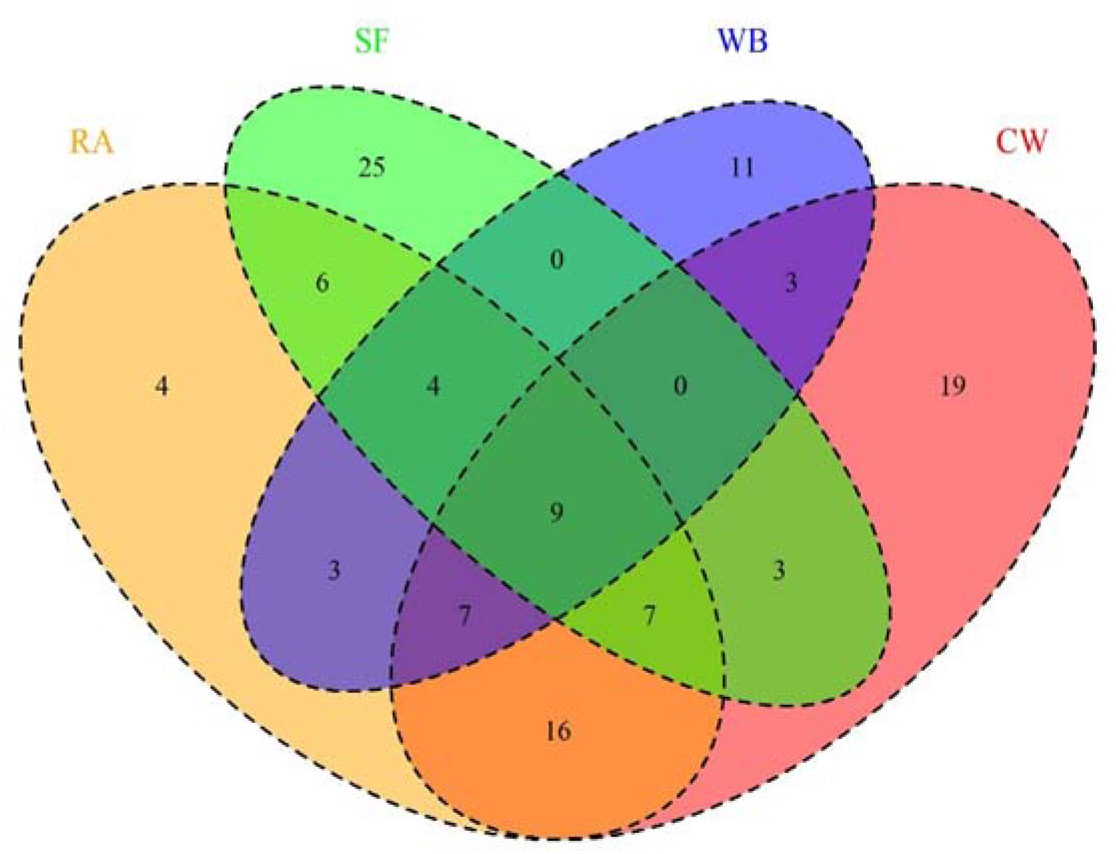
Venn diagram showing the number of unique and shared species among the different sampling habitats; CW: Cropland and Wasteland, RA: Residential Areas, SF: Sal Forest, WB: Waterbodies

### 3.4. Feeding guilds and functional diversity

Out of a total of six feeding guilds of birds observed, the richness of insectivores (49 species; 41.9%) dominates the others, followed by omnivorous (27 species; 23.1%), carnivorous (22 bird species; 18.8%), granivorous (10 bird species; 8.5%) and frugivorous (7 bird species; 6%) (Fig. 7). Only two nectarivorous feeding birds were observed. However, the individual counting of each feeding guild shows that O represents the highest numbers (2703 individuals), followed by I, C, G, F, and N, respectively. A hierarchical dendrogram cluster exhibited similarity between the guild and guild composition in our study area (Fig. 8).

**Fig. 7.**
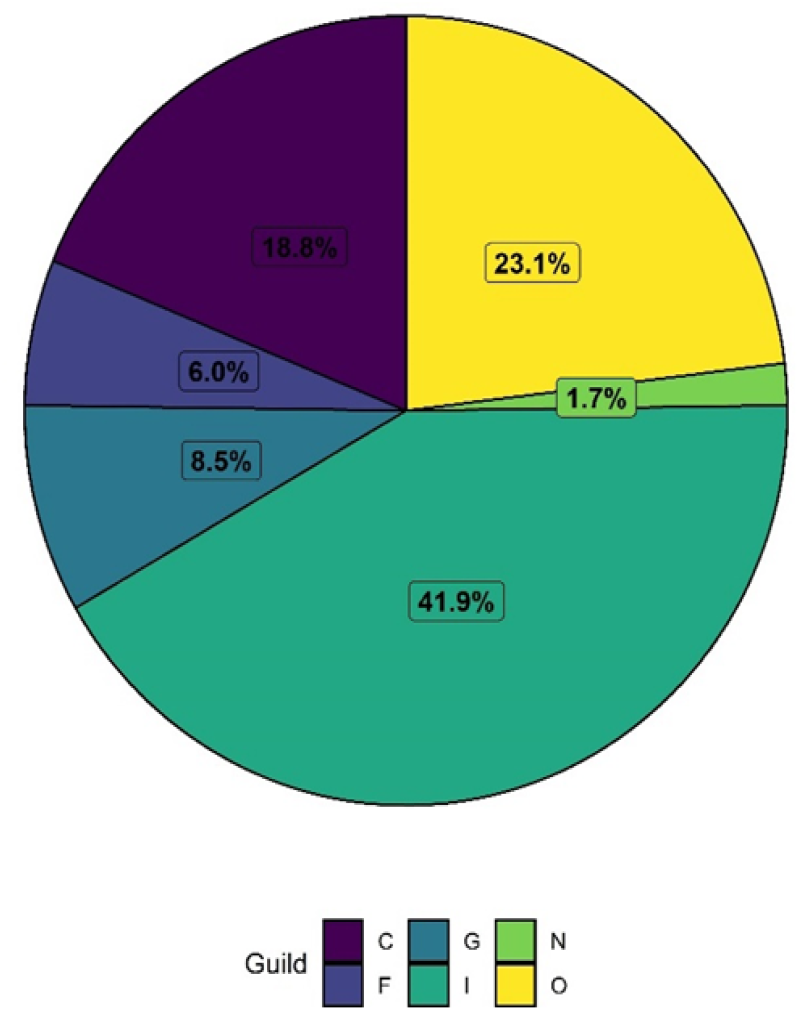
The pie chart representing the occurrence of birds feeding guild richness (%) in the study area; C= Carnivore, F = Frugivore, G = Granivore, I = Insectvore, N = Nectarivore and O = Omnivore

**Fig. 8.**
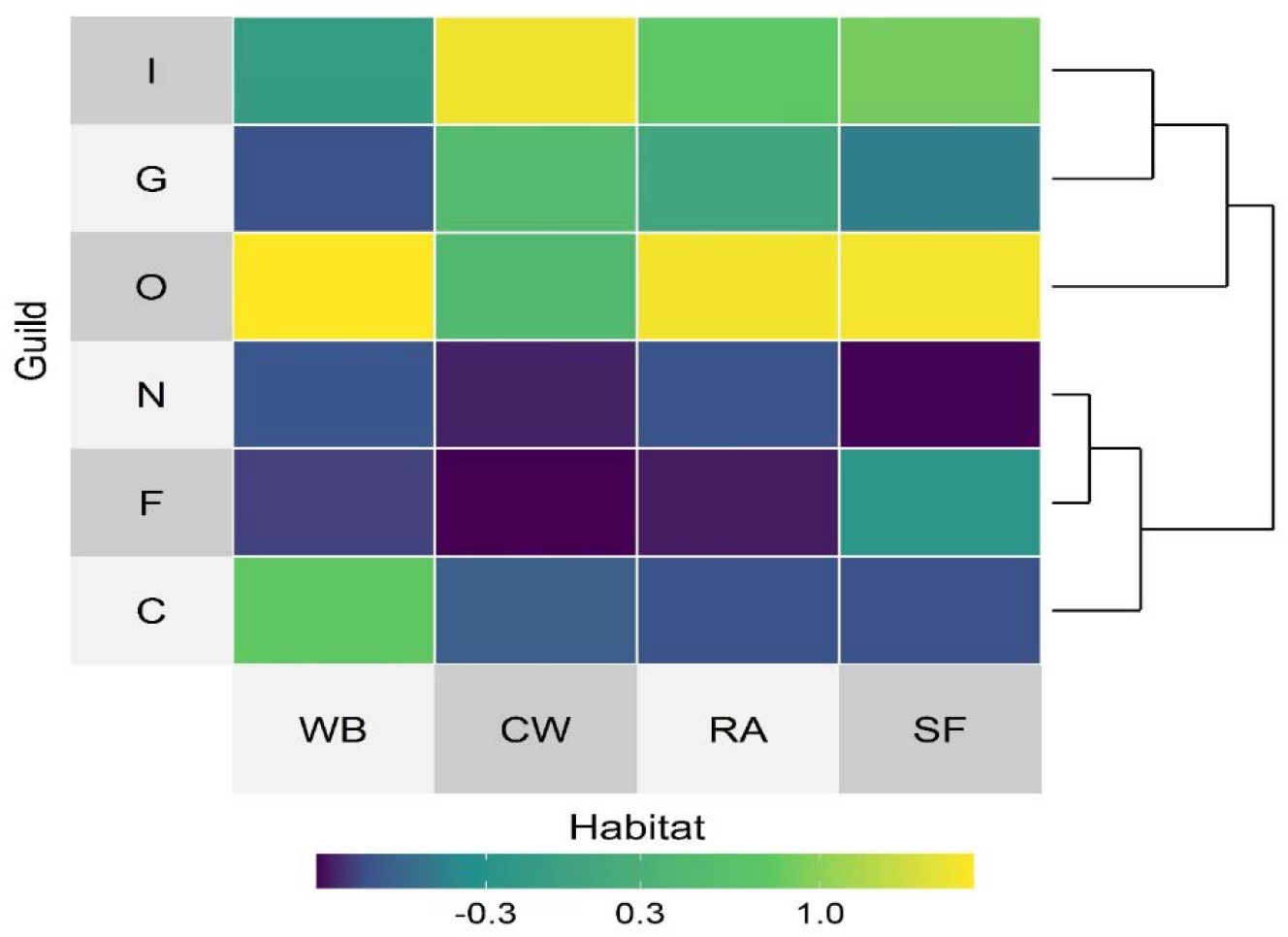
Heat map showing the distribution pattern of various feeding guild in different habitat; blue colour represented low value and yellow indicated higher value. The distribution pattern of birds’ guild in various habitat form two higherchial cluster; CW: Cropland and Wasteland, RA: Residential Areas, SF: Sal Forest, WB: Waterbodies, I: Insectvore, G: Grainivore, O: Omnivore, N: Nectarivore and C: Carnivore

## 4. Discussion

Our study demonstrated that bird richness and diversity were higher in crops and wasteland. Worldwide, it is accepted that certain bird species favour cropland habitats considering that about one-third of the world’s bird population occasionally stays in such a habitat. For example, good management of crop fields and mosaic of micro-landscape provides food resources to promote higher bird richness (Tu et al., 2020) and agricultural habitat harbouring a significant proportion of the birds’ community (Martin et al., 2012). Further, avian species shared the highest similarities among cropland and wasteland and residential areas. Abundant food sources, and several insect species attracted by the crops, may be harbouring many bird species supporting different feeding guilds (Filloy and Bellocq, 2007; Palei et al., 2012; Sánchez-Oliver et al., 2014). Residential areas shared the similarity in abundance with crop and wasteland, which implies the component of generalist birds in the study area. Mosaic of a variety of tree covers, small water bodies, bamboo grooves, and grasses around cropland area makes a more productive heterogeneous landscape that allows it to sustain various bird species (Hossain and Aditya, 2016). However, a comparison between residential areas and forest areas showed that the diversity and richness are higher in residential areas, possibly due to habitat loss in forest areas which forced the bird species to inhibit in residential areas (Sarkar et al., 2009). On the other hand, the diversity and richness of bird richness are higher in residential areas than in forest areas. In monoculture, sal-dominated forest habitats may have low fruit and flowering trees, while residential areas have parks and roadside fruit trees like banyan (*Ficus benghalensis*), peepal (*F. religiosa*), and jamun (*Syzygium cumini*), etc., in residential areas.

Water bodies allowed the highest number of birds, and their abundance revealed that water bodies significantly differed from other habitats in our study area. The abundance of birds was higher in water bodies because of the large number of gathering colonial water birds like lesser-whistling duck (*Dendrocygna javanica*), cotton pygmy duck (*Nettapus coromandelianus*), Asian openbill stork (*Anastomus oscitans*), etc. Birds are abundant in this habitat because of some environmental characteristics of wetland or water bodies like size, depth, water level, and plant species (Woldemariam et al., 2018).

The bird richness of the residential area does not exhibit any significant differences from cropland and wasteland. However, total bird counts (abundance) in this habitat significantly differ from sal forest. Certain kinds of birds were encouraged by urban resources. They are likely to be more exploitative and aggressive and can adjust to this urban environment by taking advantage of the anthropological resources (Audet et al., 2016; Charmantier et al., 2017; Senar et al., 2017).

Our study observed that the insectivorous are the richest feeding guild, followed by omnivorous, carnivorous, granivorous, frugivorous, and nectarivorous. Insectivorous species richness has been found to coincide with food availability in agricultural and residential habitats (Chatterjee et al., 2013; Gatesire et al., 2014; Mukhopadhyay and Mazumdar, 2019). The omnivorous are the second most rich-feeding guilds. Again, (Mukhopadhyay and Mazumdar (2019) explained that they have a wildly distributed guild because omnivores have an affinity to utilize natural resources and food. Our study also supported that they extend themselves in urban areas with high numbers in both spatial and temporal gradients (Clergeau et al., 1998; Jokimäki and Suhonen, 1993; Sorace, 2002). Granivorous and omnivorous tend to colonize in the degraded agricultural landscape (Frishkoff et al., 2014). However, our study explained that only granivorous showed colonization in the mosaic landscape of cropland and wasteland. In contrast, omnivores showed maximum colonization in water bodies, followed by residential and sal forest areas. The colonization of granivorous in cropland and wasteland because the open habitats provide abundant seed grain (Chettri et al., 2005; Diaz and Telleria, 1996). Fruiting plants are regulated by the frugivorous birds (Chatterjee et al., 2018; Trager and Mistry, 2003). We observed that the abundance of frugivorous birds is higher in forest areas than in residential areas because of high fruit plant diversity (Gomes et al., 2008). Nectarivores were preferably low in number compared to other guilds. They prefer open habitats and are regulated by flowering plants during the blooming season (Abrahamczyk and Kessler, 2010). We witnessed that their numbers were comparatively higher in residential areas than in other areas because nectarivorous were regulated by the blooming of banana (*F. benghalensis*), papaya (*Carica papaya*), and *Hibiscus* sp. and *Lantana camara*.

## 5. Conclusion

We observed that the diversity in the habitat regulated the birds’ diversity. Insectivorous birds were the highest feeding guilds, followed by omnivorous and granivorous. The cropland and wasteland habitat regulated bird feeding guilds and diversity. Therefore, the agricultural and degraded landscapes like cropland and wasteland played an important role in maintaining bird diversity. Trees, shrubs, and old buildings in the residential areas also provide suitable habitats for birds. However, several developmental activities like road widening, building construction, and irrigation practices altered the habitats of birds by felling old trees like *Ficus bengalensis* and *Ficus religiosa;* as a result, many frugivorous birds lost their feeding ground. Therefore, proper urban planning is required to protect bird diversity in the human-dominated landscape.

## Author contributions

HSP conceived the study. RK, AK and AG collected the data. RK and HSP performed the analyses. RK and HSP wrote the first draft of the paper. AK, AG and RKM revised the manuscript. All authors read and approved the final manuscript.

## Acknowledgements

We are thankful to the Odisha forest department and administrative authority of North Orissa University to conduct the survey. We would like to thank Mr. Russell J. Grey for his technical advice on use of R software.

## References

Abrahamczyk, S., Kessler, M., 2010. Hummingbird diversity, food niche characters, and assemblage composition along a latitudinal precipitation gradient in the Bolivian lowlands. J. Ornithol. 151, 615–625.

Ali, S., 1990. The book of Indian birds, 7th ed. Oxford University Press.

Audet, J.-N., Ducatez, S., Lefebvre, L., 2016. The town bird and the country bird: problem solving and immunocompetence vary with urbanization. Behav. Ecol. 27, 637–644.

Barth, B.J., FitzGibbon, S.I., Wilson, R.S., 2015. New urban developments that retain more remnant trees have greater bird diversity. Landsc. Urban Plan. 136, 122–129.

Beukema, H., Danielsen, F., Vincent, G., Hardiwinoto, S., van Andel, J., 2007. Plant and bird diversity in rubber agroforests in the lowlands of Sumatra, Indonesia. Agrofor. Syst. 70, 217–242.

Bibby, C.J., Burgess, N.D., Hillis, D.M., Hill, D.A., Mustoe, S., 2000. Bird census techniques, 2nd ed. Elsevier.

Blair, R.B., 1999. Birds and butterflies along an urban gradient: surrogate taxa for assessing biodiversity? Ecol. Appl. 9, 164–170.

Camacho-Cervantes, M., Ojanguren, A.F., MacGregor-Fors, I., 2018. Birds from the burgh: bird diversity and its relation with urban traits in a small town. J. Urban Ecol. 4, juy011.

Chao, A., 1984. Non-parametric estimation of the classes in a population. Scand J Stat. 11, 265–270.

Chao, A., Chazdon, R.L., Colwell, R.K., Shen, T., 2005. A new statistical approach for assessing similarity of species composition with incidence and abundance data. Ecol. Lett. 8, 148–159.

Charmantier, A., Demeyrier, V., Lambrechts, M., Perret, S., Grégoire, A., 2017. Urbanization is associated with divergence in pace-of-life in great tits. Front. Ecol. Evol. 5, 53.

Chatterjee, A., Adhikari, S., Barik, A., Mukhopadhyay, S.K., 2013. The mid-winter assemblage and diversity of bird populations at Patlakhawa Protected Forest, Coochbehar, West Bengal, India. Ring 35, 31.

Chatterjee, P., Mondal, K., Chandra, K., Tripathy, B., 2018. First photographic evidence of Asian Golden Cat Catopuma temminckii (Vigors and Horsfield, 1827) from Neora valley National Park, Central Himalayas, India. Rec. Zool. Surv. India 118, 128.

Chettri, N., Deb, D.C., Sharma, E., Jackson, R., 2005. The relationship between bird communities and habitat. Mt. Res. Dev. 25, 235–243.

Clergeau, P., Savard, J.-P.L., Mennechez, G., Falardeau, G., 1998. Bird Abundance and Diversity along an Urban-Rural Gradient: A Comparative Study between Two Cities on Different Continents. Condor 100, 413–425.

Das, U.P., Acharya, A., Palei, H.S., 2021. The diet of Indian foxes in a peri-urban area of eastern India. Acta Ecol. Sin.

Dias, S., Moreira, F., Beja, P., Carvalho, M., Gordinho, L., Reino, L., Oliveira, V., Rego, F., 2013. Landscape effects on large scale abundance patterns of turtle doves Streptopelia turtur in Portugal. Eur. J. Wildl. Res. 59, 531–541.

Diaz, M., Telleria, J.L., 1996. Granivorous birds in a stable and isolated open habitat within the Amazonian rainforest. J. Trop. Ecol. 12, 419–425.

Filloy, J., Bellocq, M.I., 2007. Patterns of bird abundance along the agricultural gradient of the Pampean region. Agric. Ecosyst. Environ. 120, 291–298.

Fontana, S., Sattler, T., Bontadina, F., Moretti, M., 2011. How to manage the urban green to improve bird diversity and community structure. Landsc. Urban Plan. 101, 278–285.

Frishkoff, L.O., Karp, D.S., M’Gonigle, L.K., Mendenhall, C.D., Zook, J., Kremen, C., Hadly, E.A., Daily, G.C., 2014. Loss of avian phylogenetic diversity in neotropical agricultural systems. Science (80-.). 345, 1343–1346.

Garden, J., McAlpine, C., Peterson, A.N.N., Jones, D., Possingham, H., 2006. Review of the ecology of Australian urban fauna: a focus on spatially explicit processes. Austral Ecol. 31, 126–148.

Gatesire, T., Nsabimana, D., Nyiramana, A., Seburanga, J.L., Mirville, M.O., 2014. Bird diversity and distribution in relation to urban landscape types in Northern Rwanda. Sci. World J. 2014.

Gomes, L.G.L., Oostra, V., Nijman, V., Cleef, A.M., Kappelle, M., 2008. Tolerance of frugivorous birds to habitat disturbance in a tropical cloud forest. Biol. Conserv. 141, 860–871.

Grimmett, R., Inskipp, C., Inskipp, T., 2016. Birds of the Indian Subcontinent: India, Pakistan, Sri Lanka, Nepal, Bhutan, Bangladesh and the Maldives. Bloomsbury Publishing.

Hamel, P.B., 1982. Bird-habitat relationships on southeastern forest lands. US Department of Agriculture, Southeastern Forest Experiment Station, Forest ….

Herzog, F., Dreier, S., Hofer, G., Marfurt, C., Schüpbach, B., Spiess, M., Walter, T., 2005. Effect of ecological compensation areas on floristic and breeding bird diversity in Swiss agricultural landscapes. Agric. Ecosyst. Environ. 108, 189–204.

Hossain, A., Aditya, G., 2016. Avian diversity in agricultural landscape: records from Burdwan, West Bengal, India. In: Proceedings of the Zoological Society. Springer, pp. 38–51.

Issa, M.A.A., 2019. Diversity and abundance of wild birds species’ in two different habitats at Sharkia Governorate, Egypt. J. Basic Appl. Zool. 80, 1–7.

Jokimäki, J., Suhonen, J., 1993. Effects of urbanization on the breeding bird species richness in Finland: a biogeographical comparison. Ornis Fenn. 70, 71–77.

Kiros, S., Afework, B., Legese, K., 2018. A preliminary study on bird diversity and abundance from Wabe fragmented forests around Gubre subcity and Wolkite town, Southwestern Ethiopia. Int. J. Avian Wildl. Biol. 3, 333–340.

Loss, S.R., Ruiz, M.O., Brawn, J.D., 2009. Relationships between avian diversity, neighborhood age, income, and environmental characteristics of an urban landscape. Biol. Conserv. 142, 2578–2585.

Magurran, A., 2004. Measuring Biological Diversity, 1st ed. Blackwell Science Ltd.

Martin, E.A., Viano, M., Ratsimisetra, L., Laloë, F., Carrière, S.M., 2012. Maintenance of bird functional diversity in a traditional agroecosystem of Madagascar. Agric. Ecosyst. Environ. 149, 1–9.

Miller, J.R., Hobbs, R.J., 2002. Conservation where people live and work. Conserv. Biol. 16, 330–337.

Mukhopadhyay, S., Mazumdar, S., 2019. Habitat-wise composition and foraging guilds of avian community in a suburban landscape of lower Gangetic plains, West Bengal, India. Biologia (Bratisl). 74, 1001–1010.

Nason, I., 1992. Discovering birds. Pisces Publication.

Niemi, G.J., 1985. Patterns of morphological evolution in bird genera of New World and Old World peatlands. Ecology 66, 1215–1228.

Palei, H.S., Mohapatra, P.P., Sahu, H.K., 2012. Birds of Hadagarh Wildlife Sanctuary, Odisha, Eastern India. World J. Zool. 7, 221–225.

Prasad, S.N., Ramachandra, T. V, Ahalya, N., Sengupta, T., Kumar, A., Tiwari, A.K., Vijayan, V.S., Vijayan, L., 2002. Conservation of wetlands of India-a review. Trop. Ecol. 43, 173–186.

Ralph, C.J., Geupel, G.R., Pyle, P., Martin, T.E., DeSante, D.F., Mila, B., 1996. Manual of field methods for monitoring terrestrial birds, US Forest Service General Technical Report PSW.

Reginald, L., Mahendran, C., Kumar, S., Pramod, P., 2007. Birds of Singanallur lake, Coimbatore, Tamil Nadu. Zoos’ Print J. 22, 2944–2948.

Sánchez-Oliver, J.S., Benayas, J.M.R., Carrascal, L.M., 2014. Local habitat and landscape influence predation of bird nests on afforested Mediterranean cropland. Acta Oecologica 58, 35–43.

Sarkar, N.J., Sultana, D., Jaman, M.F., Rahman, M.K., 2009. Diversity and population of avifauna of two urban sites in Dhaka, Bangladesh. Ecoprint An Int. J. Ecol. 16, 1–7.

Sauer, J.R., Link, W.A., Fallon, J.E., Pardieck, K.L., Ziolkowski, D.J., 2013. The North American breeding bird survey 1966–2011: summary analysis and species accounts. North Am. Fauna 1–32.

Senar, J.C., Garamszegi, L.Z., Tilgar, V., Biard, C., Moreno-Rueda, G., Salmón, P., Rivas, J.M., Sprau, P., Dingemanse, N.J., Charmantier, A., 2017. Urban great tits (Parus major) show higher distress calling and pecking rates than rural birds across Europe. Front. Ecol. Evol. 5, 163.

Sengupta, S., Mondal, M., Basu, P., 2014. Bird species assemblages across a rural urban gradient around Kolkata, India. Urban Ecosyst. 17, 585–596.

Shoffner, A., Wilson, A.M., Tang, W., Gagné, S.A., 2018. The relative effects of forest amount, forest configuration, and urban matrix quality on forest breeding birds. Sci. Rep. 8, 1–12.

Sodhi, N.S., Şekercioğlu, C.H., Barlow, J., Robinson, S.K., 2011. Effects of habitat fragmentation on tropical birds. In: Sodhi, NS, Sekercioglu, Çagan H. Barlow, J. and Robinson, S. (Eds.), Sodhi, NS, Sekercioglu, Çagan H. Barlow, J. and Robinson, S. Eds. Conservation of Tropical Birds. 1st Ed. Blackwell Publishing Ltd. p. 159.

Sorace, A., 2002. High density of bird and pest species in urban habitats and the role of predator abundance. Ornis Fenn. 79, 60–71.

Terborgh, J., 1985. The role of ecotones in the distribution of Andean birds. Ecology 66, 1237–1246.

Trager, M., Mistry, S., 2003. Avian community composition of kopjes in a heterogeneous landscape. Oecologia 135, 458–468.

Tu, H.-M., Fan, M.-W., Ko, J.C.-J., 2020. Different habitat types affect bird richness and evenness. Sci. Rep. 10, 1–10.

Verma, A., Balachandran, S., Chaturvedi, N., Patil, V., 2004. A preliminary report on the biodiversity of Mahul Creek, Mumbai, India with special reference to avifauna. Zoos’ Print J. 19, 1599–1605.

Vitousek, P.M., Mooney, H.A., Lubchenco, J., Melillo, J.M., 1997. Human domination of Earth’s ecosystems. Science (80-.). 277, 494–499.

Walther, B.A., Moore, J.L., 2005. The concepts of bias, precision and accuracy, and their use in testing the performance of species richness estimators, with a literature review of estimator performance. Ecography (Cop.). 28, 815–829.

Wang, X., Zhu, G., Ma, H., Wu, Y., Zhang, W., Zhang, Y., Li, C., de Boer, W.F., 2022. Bird communities’ responses to human-modified landscapes in the southern Anhui Mountainous Area. Avian Res. 13, 100006.

Woldemariam, W., Mekonnen, T., Morrison, K., Aticho, A., 2018. Assessment of wetland flora and avifauna species diversity in Kafa Zone, Southwestern Ethiopia. J. Asia-Pacific Biodivers. 11, 494–502.

