## Appendix A for "Bird assemblages in a peri-urban landscape in eastern India"

**Appendex A**

| Family | Scientific Name | Common Name | Code | IUCN Status | Guild | RA | WB | CW | SF |
| --- | --- | --- | --- | --- | --- | --- | --- | --- | --- |
| Psittacidae | *Psittacula eupatria* | Alexandrine parakeet | ap | NT | F | + | - | - | + |
| Alaudidae | *Eremopterix griseus* | Ashy-crown Sparrow-lark | acsl | LC | G | - | - | + | - |
| Cisticolidae | *Prinia socialis* | Ashy Prinia | asp | LC | I | - | - | + | - |
| Cuculidae | *Eudynamysscolopaceus* | Asian Koel | ak | LC | O | + | - | - | + |
| Ciconiidae | *Anastomus oscitans* | Asian Openbill Stork | aos | LC | C | - | + | + | - |
| Apodidae | *Cypsiurus balasiensis* | Asian Palm-swift | aps | LC | I | + | + | + | - |
| Monarchidae | *Terpsiphone paradisi* | Asian Paradise-flycatcher | apfc | LC | I | - | - | - | + |
| Sturnidae | *Gracupica contra* | Asian Pied Starling | apst | LC | O | + | + | + | + |
| Sturnidae | *Acridotheres ginginianus* | Bank Myna | bm | LC | O | + | - | - | - |
| Hirundinidae | *Hirundo rustica* | Barn Swallow | bs | LC | I | - | + | + | - |
| Ploceidae | *Plocus philippinus* | Baya Weaver | bw | LC | G | - | - | + | - |
| Laniidae. | *Lanius vittatus* | Bay-backed Shrike | bbs | LC | C | - | - | - | + |
| Dirucadae | *Dicrurus macrocercus* | Black Drongo | bd | LC | I | + | + | + | + |
| Estrildidae | *Lonchura antricapilla* | Black-headed Munia | bhm | LC | G | - | - | + | - |
| Accipitridae | *Milvus migrans* | Black kite | bk | LC | C | + | + | + | - |
| Accipitridae | *Elanus axillaris* | Black-winged Kite | bwk | LC | C | + | - | + | - |
| Campephagidae | *Coracina melanoptera* | Black-headed Cuckooshrike | bhcs | LC | O | - | - | - | + |
| Oriolidae | *Oriolus xanthornus* | Black-hooded Oriole | bho | LC | O | + | + | - | + |
| Oriolidae | *Oriolus chinensis* | Black-naped Oriole | bno | LC | O | - | - | - | + |
| Picidae | *Dinopium benghalense* | Black-rumped Woodpecker | brw | LC | I | - | - | - | + |
| Columbidae | *Columba livia* | Rock Pigeon | rp | LC | G | + | - | + | - |
| Megalaimidae | *Megalaima asiatica* | Blue-throated Barbet | btb | LC | F | - | - | - | + |
| Meropidae | *Nyctyornis athertoni* | Blue-bearded Bee-eater | bbbe | LC | I | - | - | - | + |
| Turnicidae | *Turnix sylvaticus* | Small Buttonquail | bq | LC | O | - | - | + | - |
| Sturnidae | *Sturnia pagodarum* | Brahminy Starling | bst | LC | O | + | - | + | + |
| Dirucadae | *Dicrurus aeneus* | Bronzed Drongo | brd | LC | I | + | - | + | + |
| Jacanidae | *Metopidius indicus* | Bronze-winged Jacana | bwj | LC | I | - | + | - | - |
| Laniidae. | *Lanius cristatus* | Brown Shrike | brs | LC | I | + | - | + | - |
| Megalaimidae | *Psilopogon zeylanicus* | Brown-headed Barbet | bhb | LC | F | - | - | - | + |
| Ardeidae | *Bubulcus ibis* | Cattle Egret | ce | LC | I | + | - | + | + |
| Meropidae | *Merops leschenaulti* | Chestnut-headed Bee-eater | chbe | LC | I | - | - | + | + |
| Sturnidae | *Sturnia malabarica* | Chestnut-tailed Starling | cts | LC | O | + | - | + | + |
| Phylloscopidae | *Phylloscopus collybita* | Common Chiffchaff | cc | LC | I | + | - | + | - |
| Rallidae | *Fulica atra* | Common Coot | cct | LC | O | - | + | - | - |
| Cuculidae | *Hierococcyx varius* | Common Hawk-Cuckoo | chc | LC | I | + | - | - | + |
| Upupidae | *Upupa epops* | Eurasian Hoopoe | eh | LC | I | - | - | + | - |
| Aegithinidae | *Aegithina tiphia* | Common Iora | ci | LC | I | - | - | - | + |
| Alcedinidae | *Alcedo atthis* | Common Kingfisher | ck | LC | C | + | + | + | - |
| Rallidae | *Gallinula chloropus* | Eurasian Moorhen | emh | LC | O | + | + | - | - |
| Sturnidae | *Acridotheres tristis* | Common Myna | cm | LC | O | + | + | + | + |
| Scolopacidae | *Actitis hypoleucos* | Common Sandpiper | csp | LC | I | - | - | + | - |
| Scolopacidae | *Gallinago gallinago* | Common Snipe | csn | LC | I | - | + | + | - |
| Cisticolidae | *Orthotomus sutorius* | Common Tailorbird | ctb | LC | I | + | - | + | - |
| Megalaimidae | *Megalaima haemacephala* | Coppersmith Barbet | csb | LC | F | + | - | - | + |
| Anatidae | *Nettapus coromandelianus* | Cotton Pygmy-Goose | cpg | LC | O | - | + | - | - |
| Accipitridae | *Spilornis cheela* | Crested Serpent-Eagle | cse | LC | C | - | - | - | + |
| Motacillidae | *Motacilla citreola* | Citrine Wagtail | cw | LC | I | + | + | + | + |
| Phylloscopidae | *Phylloscopus fuscatus* | Dusky Warbler | dw | LC | I | + | - | - | + |
| Columbidae | *Streptopelia tranquebarica* | Red Collered-Dove | rcd | LC | G | - | - | + | - |
| Columbidae | *Streptopelia decaocto* | Eurasian Collared-Dove | ecd | LC | G | - | - | + | + |
| Oriolidae | *Oriolus oriolus* | Eurasian Golden Oriole | ego | LC | O | + | + | - | + |
| Motacillidae | *Dendronanthus indicus* | Forest Wagtail | fw | LC | I | - | - | - | + |
| Picidae | *Dendrocopos macei* | Fulvous-breasted Woodpecker | fbw | LC | I | - | - | - | + |
| Cuculidae | *Centropus sinensis* | Greater Coucal | gc | LC | O | + | - | + | + |
| Picidae | *Chrysocolaptes guttacristatus* | Greater Flameback Woodpeaker | gfw | LC | I | - | - | - | + |
| Meropidae | *Merops orientalis* | Green Bee-eater | gbe | LC | I | + | + | + | + |
| Cuculidae | *Phaenicophaeus tristis* | Green-billed Malkoha | gbm | LC | C | - | - | - | + |
| Motacillidae | *Motacilla cinerea* | Grey Wagtail | gw | LC | I | + | - | + | + |
| Corvidae | *Corvus splendens* | House Crow | hc | LC | O | + | + | - | - |
| Passeridae | *Passer domesticus* | House Sparrow | hs | LC | G | + | - | + | - |
| Apodidae | *Apus nipalensis* | House Swift | hsw | LC | I | + | - | - | - |
| Caprimulg | *Caprimulgus asiaticus* | Indian Nightjar | inj | LC | I | - | - | + | - |
| Ardeidae | *Ardeola grayii* | Indian Pond-Heron | ph | LC | C | + | + | + | - |
| Muscicapidae | *Saxicoloides fulicatus* | Indian Robin | iro | LC | I | - | - | - | + |
| Coraciidae | *Coracias benghalensis* | Indian Roller | irl | LC | C | - | - | + | - |
| Cuculidae | *Clamator jacobinus* | Jacobin Cuckoo | jc | LC | I | + | - | + | - |
| Accipitridae | *Aviceda jerdoni* | Jerdon's Baza | jb | LC | C | - | - | - | + |
| Timaliidae | *Argya striata* | Jungle Babbler | jbb | LC | O | + | - | + | - |
| Sturnidae | *Acridotheres fuscus* | Jungle Myna | jm | LC | O | + | - | + | - |
| Campephagidae | *Coracina macei* | Large Cuckooshrike | jcs | LC | I | + | - | + | - |
| Ardeidae | *Ardea alba* | Large Egret | le | LC | C | - | + | - | - |
| Anatidae | *Dendrocygna javanica* | Lesser Whistling-Duck | lwd | LC | O | - | + | - | - |
| Ardeidae | *Egretta garzetta* | Little Egret | leg | LC | C | - | + | - | - |
| Podicipedidae | *Tachybaptus ruficollis* | Little Grebe | lg | LC | C | + | + | - | - |
| Muscicapidae | *Copsychus saularis* | Oriental Magpie-Robin | omr | LC | I | + | - | - | - |
| Alaudidae | *Alauda gulgula* | Oriental Skylark | osl | LC | G | - | - | + | - |
| Zosteropidae | *Zosterops palpebrosus* | Oriental White-eye | owe | LC | O | + | - | - | + |
| Motacillidae | *Anthus rufulus* | Paddyfield Pipit | pfp | LC | I | - | - | + | - |
| Jacanidae | *Hydrophasianus chirurgus* | Pheasant-tailed Jacana | ptj | LC | I | - | + | - | - |
| Alcedinidae | *Ceryle rudis* | Pied Kingfisher | pk | LC | C | - | + | - | - |
| Cisticolidae | *Prinia inornata* | Plain Prinia | pp | LC | I | + | - | + | - |
| Psittacidae | *Psittacula cyanocephala* | Pulm-headed Parakeet | php | LC | F | - | - | - | + |
| Nectariniidae | *Cinnyris asiaticus* | Purple Sunbird | psb | LC | N | + | + | + | - |
| Nectariniidae | *Leptocoma zeylonica* | Purple-rumped Sunbird | prsb | LC | N | + | + | + | - |
| Dirucadae | *Dicrurus remifer* | Lesser Racket-tailed drongo | lrtd |  | I | - | - | - | + |
| Hirundininae | *Cecropis daurica* | Red-rumped Swallow | rrsw | LC | I | - | - | + | - |
| Pycnonotidae | *Pycnonotus cafer* | Red-vented Bulbul | rvb | LC | O | + | + | + | + |
| Pycnonotidae | *Pycnonotus jocosus* | Red-whiskered Bulbul | rwb | LC | O | + | + | + | + |
| Psittacidae | *Psittacula krameri* | Rose-ringed Parakeet | rrp | LC | F | + | - | + | + |
| Sturnidae | *Pastor roseus* | Rosy Starling | rs | LC | O | + | - | + | - |
| Corvidae | *Dendrocitta vagabunda* | Rufous Treepie | rtp | LC | O | + | + | - | + |
| Picidae | *Micropternus brachyurus* | Rufous Woodpecker | rw | LC | I | - | - | - | + |
| Estrildidae | *Lonchura punctulata* | Scaly-breasted Munia | sbm | LC | G | + | - | + | - |
| Accipitridae | *Accipiter badius* | Shikra | sk | LC | C | + | + | + | + |
| Accipitridae | *Circaetus gallicus* | Short-toed Snake-Eagle | stse | LC | C | - | - | - | + |
| Columbidae | *Spilopelia chinensis* | Spotted Dove | sd | LC | G | + | + | + | + |
| Strigidae | *Athene brama* | Spotted Owlet | so | LC | C | + | + | - | + |
| Alcedinidae | *Pelargopsis capensis* | Stork-billed Kingfisher | sbk | LC | C | - | + | - | - |
| Dicaeidae | *Dicaeum agile* | Thick-billed Flowerpecker | tbfp | LC | O | - | - | - | + |
| Muscicapidae | *Ficedula albicilla* | Taiga Flycatcher | tfc | LC | I | - | - | + | - |
| Muscicapidae | *Eumyias thalassinus* | Verditer Flycatcher | vfc | LC | I | - | - | + | + |
| Rallidae | *Amaurornis phoenicurus* | White-breasted Waterhen | wbwh | LC | O | + | - | + | - |
| Alcedinidae | *Halcyon smyrnensis* | White-throated Kingfisher | wtk | LC | C | + | + | + | - |
| Motacillidae | *Motacilla alba* | White Wagtail | ww | LC | I | + | - | + | - |
| Dirucadae | *Dicrurus caerulescens* | White-bellied Drongo | wbd | LC | I | - | - | - | + |
| Artamidae | *Artamus cyanopterus* | Wood Swallow | wsw | LC | I | + | - | - | - |
| Sylviidae | *Chysomma sinensis* | Yellow-eyed Babbler | yeb | LC | I | + | - | + | - |
| Columbidae | *Treron phoenicoptera* | Yellow-footed Green-Pigeon | yfgp | LC | F | - | - | + | - |
| Cisticolidae | *Cisticola juncidis* | Zitting Cisticola | zsc | LC | I | - | - | + | - |
| Charadriidae | *Vanellus indicus* | Red-wattleed lapwing | rwl | LC | I | - | - | + | - |
| Phalacrocoracidae | *Microcarbo niger* | Little Cormorent | lc | LC | C | - | + | - | - |
| Ardeidae | *Ardea purpurea* | Purple Heron | phr | LC | C | - | + | - | - |
| Chloropseidae | *Chloropsis jerdoni* | Jerdon's Leafbird | jlb |  | I | - | - | - | + |
| Campephagidae | *Pericrocotus speciosus* | Scarlet Minivet | sm | LC | I | - | - | - | + |
| Phasianidae | *Ortygornis pondicerianus* | Grey Francolin | gf | LC | O | - | - | + | - |
| Pittidae | *Pitta brachyura* | Indian Pitta | ip | LC | I | - | - | - | + |
| Threskiornithidae | *Plegadis falcinellus* | Glossy Ibis | gi | LC | C | - | - | + | - |
